# Light-mediated decreases in cyclic di-GMP levels are potentiated by pyocyanin and inhibit structure formation in *Pseudomonas aeruginosa* biofilms

**DOI:** 10.1101/797357

**Authors:** Lisa Juliane Kahl, Alexa Price-Whelan, Lars E. P. Dietrich

## Abstract

Light is known to trigger regulatory responses in diverse organisms including slime molds, animals, plants, and phototrophic bacteria. However, light-dependent processes in non-phototrophic bacteria, and those of pathogens in particular, have received comparatively little research attention. In this study, we examined the impact of light on multicellular development in *Pseudomonas aeruginosa*, a leading cause of biofilm-based bacterial infections, using a colony morphology assay. In this assay, *P. aeruginosa* strain PA14 grown in the dark forms vertical structures (i.e., “wrinkles”) on the third day of incubation. We found that growth in blue light inhibited wrinkle formation until the fifth day and that this required the phenazine pyocyanin, a redox-active metabolite produced by PA14. Light-dependent inhibition of wrinkling was also correlated with low levels of cyclic di-GMP (c-di-GMP), consistent with the role of this signal in stimulating biofilm matrix production. Though phenazine-null biofilms also showed lower levels of c-di-GMP and subtle effects on wrinkling when grown in the light, their overall levels of c-di-GMP were higher than those of the wild type. This indicates that phenazines and light simultaneously promote c-di-GMP degradation such that c-di-GMP is pushed to a minimum level, yielding a pronounced macroscopic phenotype. A screen of enzymes with the potential to catalyze c-di-GMP synthesis or degradation identified four proteins that contribute to light-dependent inhibition of biofilm wrinkling. Together, these results provide a foundation for understanding the significance of light-dependent regulation in *P. aeruginosa*.

**Importance:** The response to light in non-phototrophic bacteria (i.e., chemotrophs) is relatively understudied in comparison to light-mediated behavior in eukaryotes and phototrophic bacteria. Though they do not depend on light for growth, chemotrophic bacteria could benefit from sensing this cue when it correlates with other parameters that are important for metabolism. In this paper, we describe light-dependent effects on a cellular signal that controls the development of multicellular assemblages, called biofilms, in *Pseudomonas aeruginosa*. We found that light at intensities that are not harmful to human cells inhibited biofilm maturation. *P. aeruginosa* is a leading cause of chronic lung infections in people with cystic fibrosis and of hospital-acquired infections. As *P. aeruginosa*’s recalcitrance to treatment is attributed in part to its facile formation of biofilms, this study provides insight into a mechanism that could be inhibited via new therapeutic tools, such as photodynamic therapy.

## Introduction

Most organisms experience some degree of light exposure, which is habitat-specific and temporally affected by Earth’s rotation. Light quality and intensity directly influence the growth of phototrophic bacteria, which are known to regulate gene expression in response to these parameters. However, light can also affect regulation in chemotrophic bacteria, serving as a proxy for correlating conditions such as those found inside animal hosts or at specific depths in freshwater environments (1). While light-dependent regulation has been described in detail for model phototrophs (1), similar mechanisms that may function in bacteria that do not use light as a source of energy are understudied.

Some investigation of photosensitivity in chemotrophs has been prompted by the relatively recent identification of light-sensing proteins in diverse non-phototrophic bacteria. In these photosensory proteins, light alters the chemical properties of cofactors, such as flavin or bilin derivatives, and changes in cofactor domains influence interactions with effector domains or other proteins with regulatory functions (2). Such interactions can induce global changes in gene expression and modulate bacterial development, sociality, and behavior. In the plant pathogen *Xanthomonas campestris*, for example, red light inhibits exopolysaccharide excretion, biofilm development, and virulence via a bacteriophytochrome that binds a bilin cofactor (3). In *Caulobacter crescentus*, *Bacillus subtilis*, and *Brucella abortus*, photosensory proteins link the effects of blue light on FMN cofactors to regulation of adhesion, the general stress response, and pathogenicity (4–6). Furthermore, in both chemotrophs and phototrophs, photosensory domains are often found in proteins that are predicted to bind, synthesize, or degrade the signaling molecule cyclic di-GMP (c-di-GMP), underscoring the potential for light to affect multicellularity and motility (7–11).

The formation of biofilms, multicellular structures held together by an excreted matrix, contributes to virulence in the hospital-acquired pathogen *Pseudomonas aeruginosa.* A major goal of research in our laboratory is to define the physiological responses of *P. aeruginosa* biofilms to different growth conditions and the traits that are relevant for host colonization. These studies have led us to examine the metabolism of phenazines, compounds excreted by *P. aeruginosa* that shuttle electrons from cells in hypoxic biofilm subzones to oxygen available closer to the biofilm periphery (12). Phenazine production balances the intracellular redox state and influences biofilm morphogenesis by inhibiting matrix production (13–15). We have also shown that biofilm-specific phenazine synthesis is required for full virulence in a mouse lung infection model (16).

In the study described here, we investigated the effects of another environmental cue— light exposure—on biofilm development in *P. aeruginosa* and found that specific wavelengths of light inhibited biofilm structure development. Our results indicate that light, like phenazines, modulates biofilm formation via c-di-GMP-dependent mechanisms. We propose a model for the integrated roles of cellular redox state and c-di-GMP in the response of *P. aeruginosa* biofilms to light.

## Results & Discussion

### Light delays wrinkling in *P. aeruginosa* PA14 colony biofilms

We use a standardized colony morphology assay to study the physiology of biofilm development in *P. aeruginosa* PA14 (15, 17). In this assay, liquid-culture aliquots are spotted onto an agar-solidified medium containing tryptone and the dyes Congo red and Coomassie blue, and incubated under controlled conditions in the dark. On the third day of growth, i.e., at ~66-72 hours, wild-type PA14 biofilms form a circular ridge at the boundary of the original culture droplet. Over the next 24-36 hours, a whorled wrinkle pattern forms inside this ridge, and short spoke-like wrinkles form that emanate from the ridge toward the edge of the colony (Fig. 1A, **top; Movie S1 & S2**) (15). To test the effect of light on PA14 biofilm development, we grew wild-type biofilms under white broad-spectrum light at ~16 µmol photons m^−2^ s^−1^ or ~3.7 W m^−2^, an intensity that is roughly an order of magnitude lower than that of sunlight on a winter day in Northern Europe (18). We found that biofilm wrinkling was delayed by ~60 hours under these conditions (Fig. 1A, **top; Movie S3 & S4**) and that the biofilms appeared to bind less Congo red, suggesting that they produced less matrix (17).

**Figure 1.**
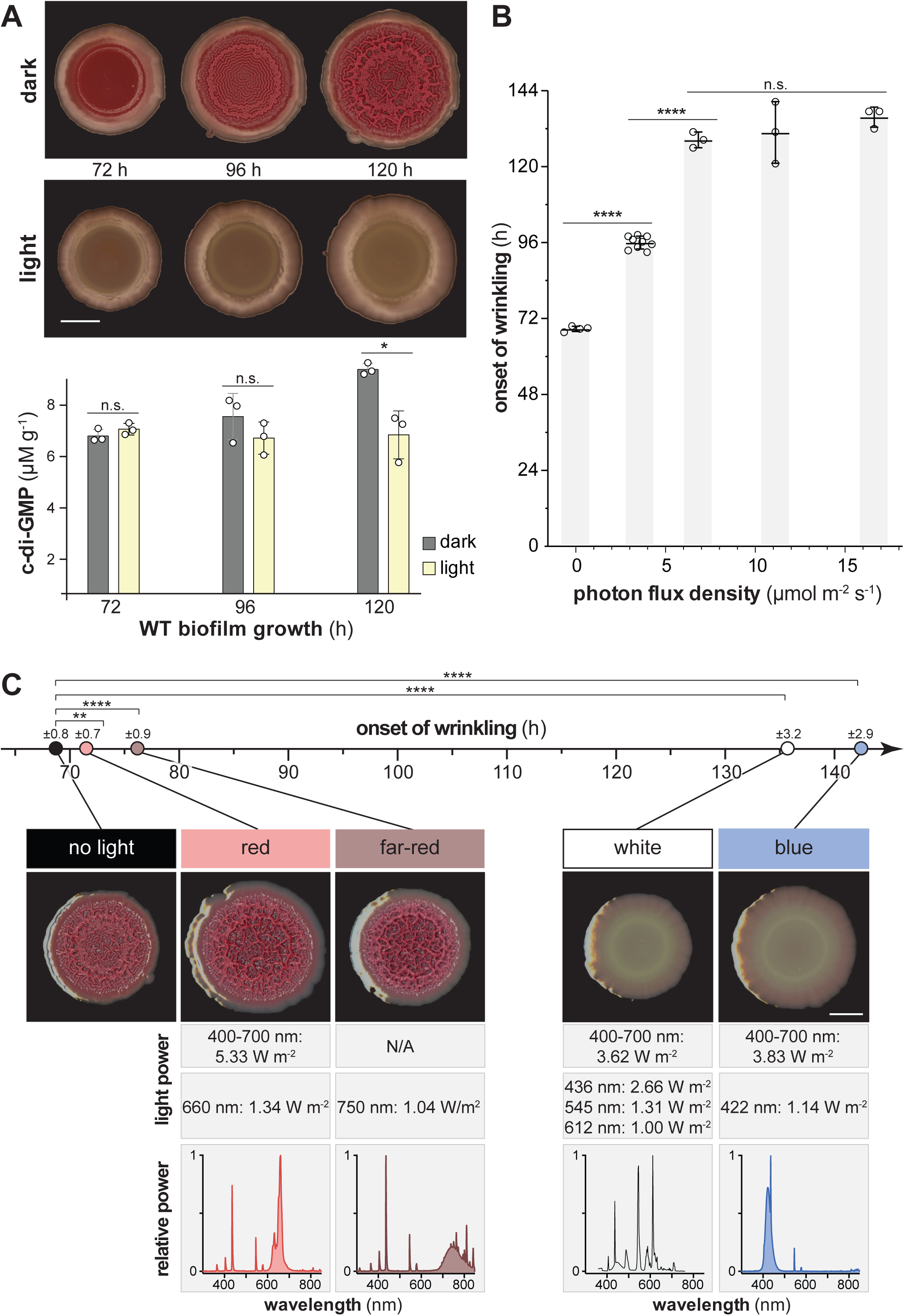
Blue light delays the onset of vertical structure (wrinkle) formation in *P. aeruginosa* PA14 colony biofilms. Biofilms were grown on colony morphology assay medium in various light conditions. (A) Top: WT biofilms grown in the dark or under white fluorescent light at ~16.5 µmol photons m^−2^ s^−1^. Representative biofilms for at least three independent experiments are shown. Scale bar is 5 mm. Bottom: c-di-GMP extracted from biofilms grown in the dark or light and normalized to dried biomass. Each bar represents the mean of three biological replicates (circles) and error bars show standard deviation; \**p* ≤ 0.05, n.s. = not significant. (B) Onset of wrinkling for WT biofilms grown in white fluorescent light at different intensities. Light intensity was measured as photon flux density between 400 nm and 700 nm. Each bar shows the mean of three biological replicates (circles) and error bars show standard deviation; \*\*\*\**p* ≤ 0.0001, n.s. = not significant. (C) Onset of wrinkling for WT biofilms grown in the dark or under the light sources indicated (relative spectral power distribution shown in bottom row). Images are representative of the 120-h time point and scale bar is 5 mm. Means of at least three biological replicates per light source are represented by circles (timeline, top). Standard deviation is listed atop each circle; \*\**p* ≤ 0.01, \*\*\*\**p* ≤ 0.0001. Light power values for the range between 400 nm and 700 nm and at the determining wavelengths are listed below each biofilm panel.

*P. aeruginosa* PA14 biofilm wrinkling is a consequence of matrix excretion, which is stimulated by the cellular signal c-di-GMP (19, 20). To examine whether the effect of light on biofilm structure formation might be mediated by c-di-GMP, we measured c-di-GMP levels in biofilms grown in the presence and absence of broad-spectrum white light (Fig. 1A, **bottom**). Biofilms grown under light showed lower c-di-GMP levels than those grown in the dark, suggesting that the effect of light exposure on biofilm development is mediated by pathways that control c-di-GMP levels.

Next, we tested whether the intensity or quality of light affected the onset of biofilm wrinkling by subjecting biofilms to a range of intensities of white broad-spectrum light, or growing them under distinct light spectra, and carrying out time-lapse imaging. Biofilms grown at a white broad-spectrum light intensity of ~3.5 µmol photons m^−2^ s^−1^ showed a delayed, intermediate onset of wrinkling, while most of those grown at intensities of ~7 µmol photons m^−2^ s^−1^ and higher did not wrinkle until the end of the experimental timeframe at ~130 hours (Fig. 1B). Biofilms grown under blue light showed a pronounced inhibition of wrinkling, while those grown under red or far-red light showed minor delays in onset of wrinkling (Fig. 1C). Together, these results support a model in which blue light promotes c-di-GMP degradation and/or inhibits c-di-GMP synthesis, thereby inhibiting matrix production and promoting biofilm smoothness.

### The phenazine pyocyanin is required for pronounced inhibition of biofilm wrinkling in the presence of light

Phenazine production is a key factor that determines the morphogenetic development of PA14 colony biofilms (21). In *P. aeruginosa*, the phenazine biosynthetic pathway yields phenazine-1-carboxylic acid (PCA), phenazine-1-carboxamide (PCN) and pyocyanin (PYO) (Fig. 2A). 5-methylphenazine-1-carboxylic acid (5-Me-PCA) is an unstable intermediate between PCA and PYO that can be non-enzymatically converted to other methylated derivatives, such as aeruginosins, under distinct conditions (21–27). When grown in the dark, phenazine-null (∆*phz*) biofilms show a much earlier onset of wrinkling, at ~40 hours of incubation, than those formed by the wild type (Fig. 2B; **Movie S1 & S2**)(15, 28). We have previously shown that the natural phenazines 5-Me-PCA and PYO and the synthetic phenazine PMS are all sufficient to inhibit PA14 biofilm wrinkle formation in the absence of light (21, 28). Therefore, it is specifically methylated phenazines that inhibit wrinkling in the dark.

**Figure 2.**
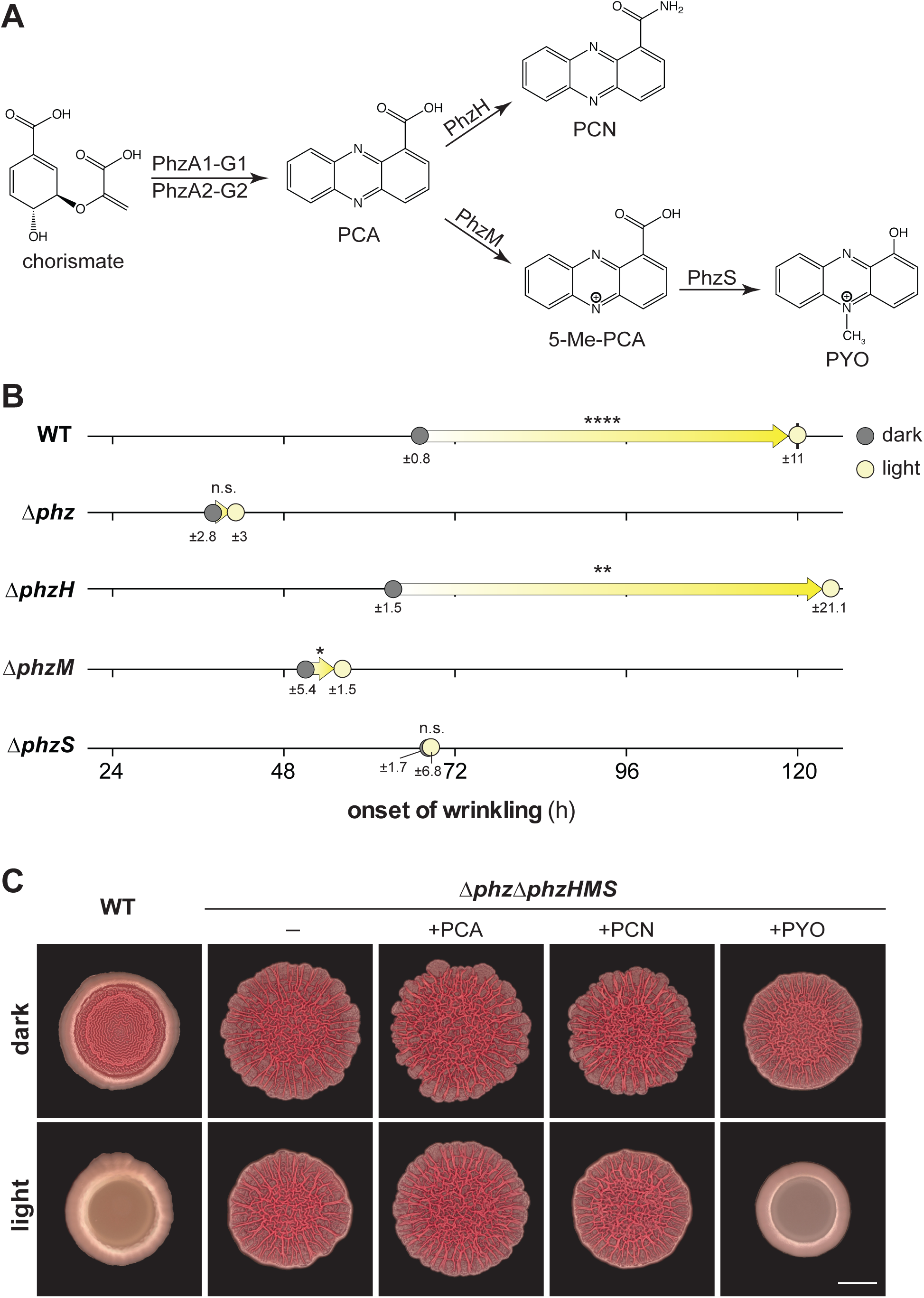
Pyocyanin (PYO) is required for pronounced inhibition of colony biofilm wrinkling in the light. (A) The biosynthetic pathway for phenazine production in *P. aeruginosa*. PCA: phenazine-1-carboxylic acid; PCN, phenazine-1-carboxamide; 5-Me-PCA, 5-methyl-phenazine-1-carboxylic acid; PYO, pyocyanin. (B) Onset of wrinkling for colony biofilms of PA14 WT and the indicated mutants when grown on colony morphology assay medium. Circles represent average in onset of wrinkling for at least three biological replicates and standard deviation is listed below each circle. Arrows designate difference in onset of wrinkling in the light compared to dark. *p*-values were determined by comparing onset of wrinkling for each mutant in the light vs. the dark and are listed atop each arrow; \**p* ≤ 0.05, \*\**p* ≤ 0.01, \*\*\*\**p* ≤ 0.0001, n.s. = not significant. (C) Biofilms of PA14 WT or the ∆*phz*∆*phzHMS* mutant grown on colony morphology assay medium amended with 200 µM of the phenazines PCA, PCN, or PYO as indicated. Images shown were taken at the 96 h time point. Scale bar is 5 mm.

Because specific phenazines critically affect the morphogenesis of PA14 biofilms grown in the absence of light, we sought to test the hypothesis that they contribute to the light-dependent effects on biofilm development. We grew phenazine biosynthetic mutants in the presence and absence of light and monitored the onset of biofilm wrinkling using time-lapse imaging. We observed that pronounced light-dependent inhibition of wrinkling required PYO production (Fig. 2B; **Movies S3 & S4**). The ∆*phzS* mutant showed no effect of light on the onset of wrinkling, which indicates that 5-Me-PCA and its derivatives other than PYO are not sufficient to mediate the light-dependent delay in wrinkling seen in the wild type.

To test whether PYO alone is sufficient to mediate light-dependent inhibition of biofilm wrinkling, we grew a ∆*phz∆phzHMS* mutant, which is not able to produce or modify phenazines, on medium containing either of the exogenously provided phenazines PCA, PCN, or PYO. We found that PYO is sufficient to inhibit wrinkling in the presence of light (Fig. 2C). ∆*phz∆phzHMS* biofilms grown under all other conditions, including those grown in the absence of phenazines, showed minor, reproducible effects of light on colony morphology; most notably, light appeared to cause a “smoothing” of the outer rim of the biofilm. However, none of these effects were as dramatic as that of PYO. These results suggest that, although 5-Me-PCA production is sufficient to delay the onset of wrinkling in biofilms grown in the dark, the delay caused by light is specifically mediated by PYO. In addition to their differences in stability and redox potential, 5-Me-PCA and PYO differ in charge and hydrophobicity, properties that may affect their reactivity with bound cofactors such as flavins and that may account for their differential roles in dark- and light-grown biofilms (21).

### Growth in the light does not affect PYO production or redox state

Prior work from our group has shown that, in the absence of light, oxidized phenazines and/or their effects on the cellular redox state promote c-di-GMP degradation and thereby inhibit biofilm wrinkling (13). We therefore wondered whether an increased abundance or a more oxidized redox state of phenazines could be responsible for the effect of light on PA14 biofilm morphogenesis. We measured phenazine production by wild-type PA14 biofilms grown in the presence and absence of light and found that light exposure did not enhance phenazine production (Fig. 3A); thus, the effect of light on PA14 colony development cannot be attributed to an increase in phenazine concentration.

**Figure 3.**
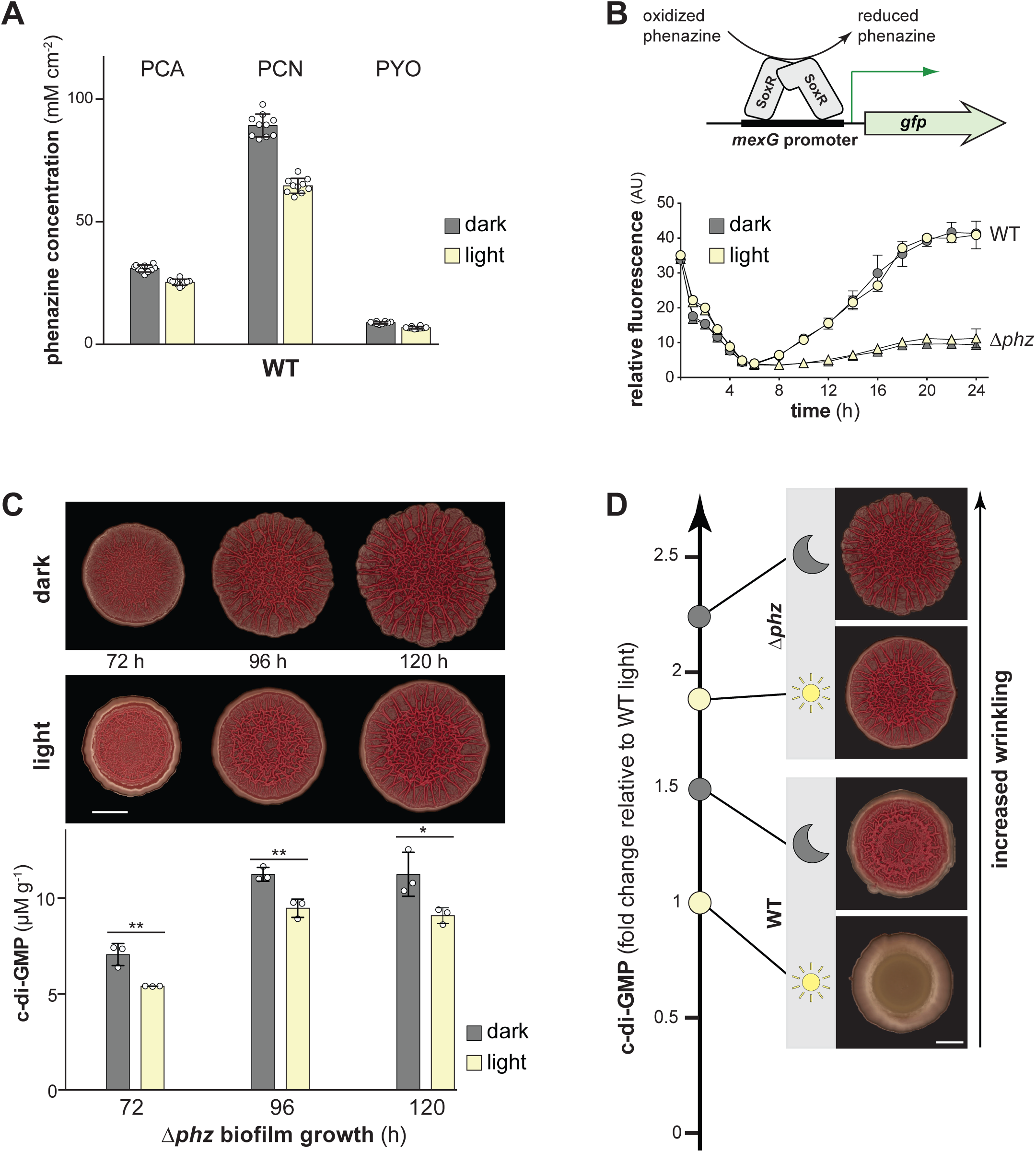
Light does not increase phenazine production or oxidize the phenazine pool, but does decrease c-di-GMP levels in the absence of phenazines. (A) Production of PCA, PCN, and PYO by colony biofilms grown in the dark or light for 96 h. Phenazines were extracted from both the biofilm and the underlying agar for each sample. Phenazine concentration was normalized to colony area. Bars represent the means of 10 biological replicates (circles) and error bars show standard deviation. (B) Top: Schematic showing the mechanism by which SoxR drives expression from the *mexG* promoter. Phenazines oxidize SoxR and trigger a conformational change that is transduced to the DNA and promotes transcription. Bottom: Phenazine- and light-dependent expression of the SoxR-regulated *mexG* reporter in PA14 WT and ∆*phz*. Expression of *attB*::P*mexG*-*gfp* in WT (circles) and ∆*phz* (triangles) was determined by measuring GFP fluorescence (excitation: 480 nm; emission: 510 nm) and normalized to the corresponding optical density at 500 nm. Data points represent averages of three biological replicates, error bars represent standard deviation. (C) Top: PA14 ∆*phz* biofilms grown in the dark or under white fluorescent light at ~16.5 µmol photons m^−2^ s^−1^. Experiment was repeated at least three times. Scale bar is 5 mm. Bottom: c-di-GMP from biofilms grown on colony morphology assay medium and normalized to dried biomass. Bars indicate the averages of three biological replicates (circles) and error bars show standard deviation; \**p* ≤ 0.05, \*\**p* ≤ 0.01. (D) Relative c-di-GMP levels of WT and ∆*phz* biofilms grown in dark and light at the 120-h time point, normalized to levels for WT grown in light. Scale bar is 5 mm.

To test whether light exposure affects the redox state of phenazines in PA14 biofilms, we employed reporter strains in which *gfp* expression is controlled by the promoter found upstream of *mexG*. The *mexGHI-opmD* operon is driven by SoxR, a redox-sensitive transcription factor that is directly activated by oxidized phenazines (29–31). When we grew P*mexG-gfp* reporter strains in liquid cultures, we found that growth in the presence of light had no effect on fluorescence (Fig. 3B). The ∆*phz* control strains showed substantially lower levels of fluorescence, consistent with the role of phenazines in activating SoxR and *mexGHI-opmD* expression. We conclude that light exposure does not affect the redox state of phenazines in PA14.

### Biofilms grown with light exposure show PYO-independent lowering of c-di-GMP levels

Prior work from our group has linked oxidized phenazines to c-di-GMP degradation and inhibition of biofilm wrinkling (13). Our observations that c-di-GMP levels are lower in light-exposed PA14 biofilms and that PYO is the primary determinant of light-dependent inhibition of wrinkling raised the question of whether PYO is required for the effects of light on c-di-GMP levels. c-di-GMP measurements performed on ∆*phz* biofilms grown in the dark and the light showed that overall c-di-GMP levels were higher for both conditions than they were for wild-type biofilms grown under either condition (Fig. 3C, **bottom and** D). However, they also revealed that c-di-GMP levels were lower in light-exposed than in dark-grown ∆*phz* biofilms, indicating a phenazine-independent mechanism that either inhibits c-di-GMP synthesis or stimulates c-di-GMP degradation in response to light. The subtle effect of light on ∆*phz* biofilm morphogenesis, i.e., slightly less wrinkle formation and smoothing of the biofilm edge (Figure 3C, **top; Movie S1-S4**), may be attributable to this activity. Overall, these results suggest that the pronounced inhibition of wrinkling seen in the presence of light and PYO occurs when c-di-GMP levels are critically low, as depicted in the model in Fig. 3D. We infer that light-and phenazine-sensing are integrated, within the *P. aeruginosa* regulatory network, at the point of c-di-GMP synthesis or degradation activities and thereby regulate biofilm formation.

### Phosphodiesterases (PDEs) contribute to the light-dependent delay in biofilm structure formation

Cellular c-di-GMP levels are modulated by diguanylate cyclases (DGCs), which synthesize c-di-GMP, and phosphodiesterases (PDEs), which degrade it (32). The *P. aeruginosa* genome encodes 40 proteins with domains implicated in DGC or PDE activity (13, 33, 34). To identify proteins that could link light sensing to modulation of c-di-GMP levels and therefore biofilm matrix production (Fig. 4A), we screened mutants representing each of these proteins for light-dependent inhibition of biofilm wrinkling. We found four mutants that showed attenuated responses to light when compared to the wild type (Fig. 4B; **Movie S5-S8**). All four of the proteins represented by these mutants contain tandem GGDEF-EAL domains. GGDEF domains have the potential for DGC activity, while EAL domains have the potential for PDE activity. Proteins with tandem GGDEF-EAL domain arrangement typically exhibit PDE activity *in vivo* because their GGDEF domains often contain a divergent motif that is catalytically inactive (35).

**Figure 4.**
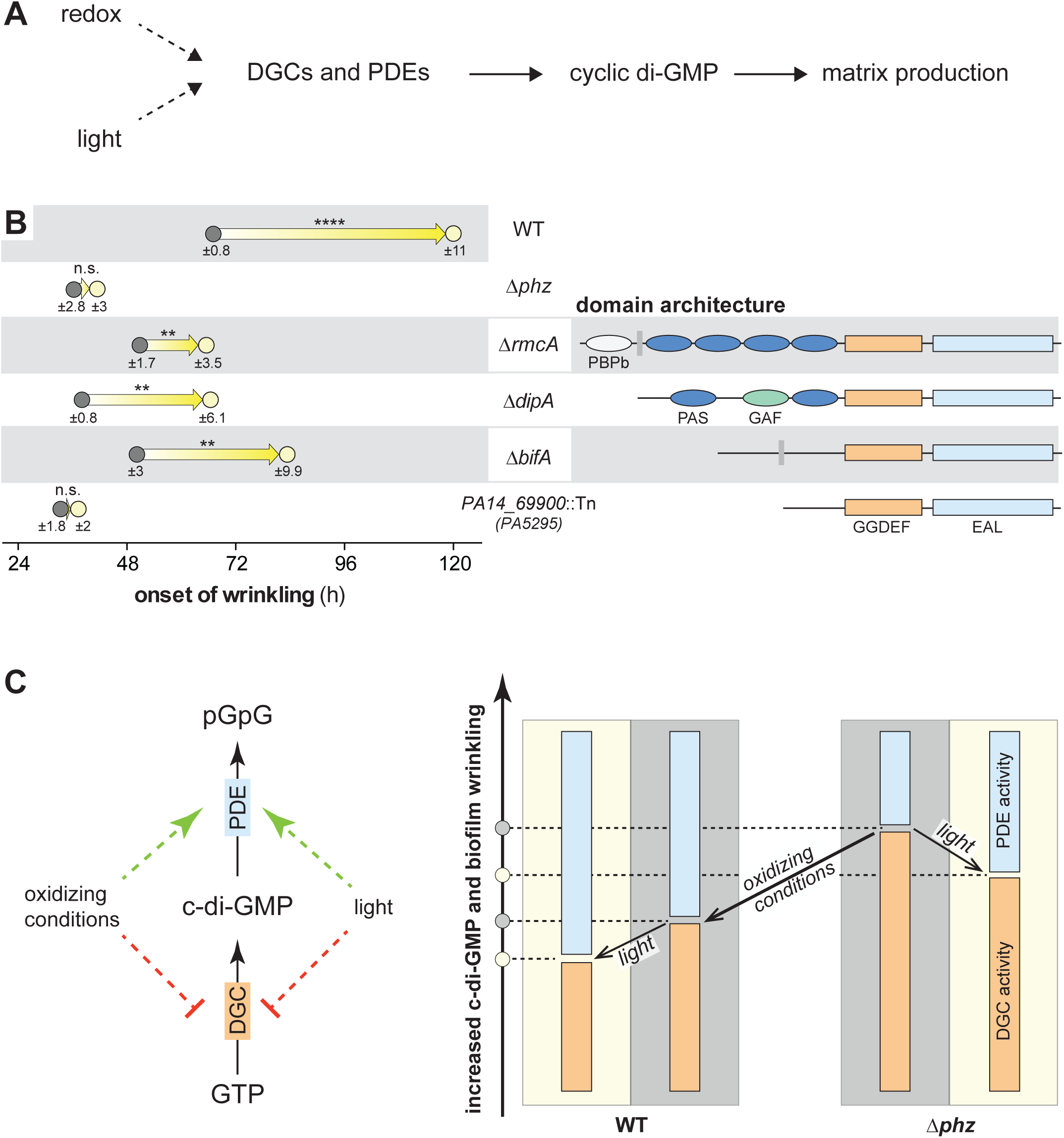
c-di-GMP levels are altered in dark vs light through dedicated sensing mechanisms. (A) Schematic for proposed influence of external inputs, redox and light, on c-di-GMP levels and matrix production via c-di-GMP modulating enzymes containing diguanylate cyclase (DGC) and phosphodiesterase (PDE) domains. (B) Left: Onset of wrinkling for colony biofilms of PA14 WT and the indicated mutants grown on colony morphology assay medium. Circles represent mean for at least three biological replicates and standard deviation is listed below each circle. Arrows designate difference in onset of wrinkling in the light compared to dark. *p*-values were determined by comparing onset of wrinkling for each mutant in the light vs. the dark and are listed atop each arrow; \*\**p* ≤ 0.01, \*\*\*\**p* ≤ 0.0001, n.s. = not significant. Right: Domain architecture of c-di-GMP modulating enzymes. Blue ellipses represent PAS domains, green ellipses represent GAF domains, grey ellipses represent PBPb domains. DGC and PDE domains are designated by orange and light blue rectangles, respectively. (C) Left: Schematic of how external factors, i.e., oxidizing conditions or light, influence c-di-GMP modulating enzymes, specifically DGC and PDE activity. Right: Model shows how c-di-GMP levels decrease with oxidizing conditions, i.e., absence of phenazines, and under light exposure in both PA14 WT and ∆*phz* colony biofilms.

The first protein represented by our screen hits, RmcA, is an exception among GGDEF-EAL domain proteins because its GGDEF and EAL motifs are both intact. In previous studies examining the role of RmcA in sensing phenazines and inhibiting biofilm wrinkling in the dark, we obtained evidence suggesting that this protein functions predominantly as a PDE but that it has DGC activity that could operate under distinct conditions (13). RmcA also contains four Per-Arnt-Sim (PAS) domains at the N-terminus. PAS domains are broadly distributed throughout the tree of life and are frequently involved in sensing stimuli (34, 36). Some of the best-known examples of PAS domain-containing proteins mediate behavioral responses to changes in light exposure, oxygen availability, or the cellular redox state (1, 8, 37–40). Based on sequence analysis and modeling, we have suggested that the four PAS domains, in order from the N-terminus, could bind 1) phenazine, 2) lipid, 3) heme, and 4) FAD, respectively (13, 34). The capacity for heme and FAD binding is of interest because porphyrins and flavins are critical elements of photosensing in diverse systems (2). Our group has also previously reported an *in vitro* characterization of RmcA’s ability to bind phenazines, which (based on a one-site binding model) indicated binding affinities (K_d_’s) of 3 µM, 6 µM, and 16 µM for PYO, phenazine methosulfate, and PCA, respectively. In this study, RmcA-deficient mutants showed a marked attenuation of the effect of light when compared to wild type. We also noted that these mutants showed less of an effect, qualitatively, on the overall pattern of wrinkling in response to light when compared to the wild type and other screen hits (**Movies S5 & S7**). Together, these observations suggest RmcA as one of the primary mediators linking photosensing to c-di-GMP degradation in *P. aeruginosa*.

Our screen identified three additional proteins with tandem GGDEF-EAL domains. In each of these proteins, the GGDEF motif is not intact, indicating that they function as PDEs *in vivo* which is consistent with published observations (13, 35, 41, 42). DipA contains an N-terminal PAS-GAF-PAS domain architecture reminiscent of the PAS-GAF-PHY arrangement that has been described for phytochromes that act as light-sensing regulatory proteins in diverse phototrophs (43). BifA and PA14_69900 are not known to contain sensory domains and may interact with other photosensory proteins that modulate their activities.

#### Concluding remarks

The results of this study suggest that integrated information regarding redox and light conditions modulates wrinkle development in PA14 biofilms. We observed that exposure to phenazines, which oxidizes the cellular redox state, and blue light exposure lead to lower c-di-GMP levels (Fig. 4C, **left panel**) and that their cumulative effect profoundly delays the onset of biofilm structure formation (Fig. 4C). In flavoproteins that respond to blue light, high photosensitivity requires that the cofactor be in its oxidized state (44, 45). We speculate that the chemical properties of PYO specifically allow it to modulate the redox states of cellular flavoproteins such that the majority are present in their oxidized forms and that those with the capacity for light sensing therefore show enhanced activity. Electron transfer to PYO has been demonstrated for the flavoprotein dihydrolipoamide dehydrogenase (46). In addition, synthetic phenazines were employed and necessary in experiments demonstrating that the cytoplasmic redox potential can shift the photosensitivity of the light-sensing kinase LovK from *C. crescentus* (47, 48). It is intriguing that PYO was unique among the *P. aeruginosa* phenazines in its capacity to potentiate light-dependent effects on biofilm morphogenesis (Fig. 2C). This finding, in combination with the fact that an RmcA-deficient mutant showed marked attenuations in these effects (Fig. 2C and **Movies S5 & S7**) and that PYO showed the highest affinity for RmcA in previous binding studies (13), suggests the potential for RmcA to sense light in a PYO-dependent manner. We speculate that RmcA integrates phenazine-dependent light and redox signals and transduces this information into regulation of biofilm development in PA14.

In addition to revealing the critical role of PYO in the effect of light on PA14 biofilm development, our observations suggest that this inhibition of wrinkling is mediated via the cellular signal c-di-GMP. Figure 4C (**right-hand panel**) illustrates the relationships between light, redox conditions, and c-di-GMP levels we observed for WT and ∆*phz* biofilms in this study. Oxidation of the cellular redox state, a consequence of phenazine production (14, 15), accounts for a larger relative decrease in c-di-GMP levels than light exposure. Nevertheless, the combined effects of phenazine production and light exposure yield a remarkably low level of c-di-GMP that is manifested as an extreme delay in biofilm structure formation. These results show that *P. aeruginosa* integrates distinct environmental and physiological cues to regulate multicellular behavior. We note that our c-di-GMP measurements were performed on extracts from whole biofilms, meaning that differences in c-di-GMP levels between subpopulations within biofilms could not be discerned. This may explain why changes in c-di-GMP levels were most consistent with effects on morphology in mature biofilms, but less so in early biofilms (i.e., compare Fig. 1A and 3C).

Recently, studies of photodynamic therapy (PDT) have revealed light, simultaneously applied with antibiotics, as a potentiator of biofilm infection eradication (49–51). Our finding that PYO, also classified as an antibiotic (52), in conjunction with light diminishes biofilm integrity is consistent with the notion that an oxidizing environment, which is often created as a result of antibiotic treatment, and light together enhance remediation of biofilm-based infections. From the perspective of bacterial physiology, changes in light intensity can accompany changes in other environmental parameters, such as nutrient availability, and chemotrophs could therefore benefit from using light as a proxy signal for conditions that directly affect their own survival. Furthermore, because eukaryotic host organisms show physiological fluctuations that are tuned to light/dark cycling, it makes sense for pathogenic bacteria to monitor light cues that correlate with these changes, for example variation in immune responses depending on the time of day (53). Finally, a change in light intensity has been speculated to serve as a proxy signal for entry into the host, allowing pathogens to trigger mechanisms of colonization and virulence appropriately (54–57). For these reasons, and because biofilm formation and maintenance are critical components of many types of infections caused by *P. aeruginosa*, the identification of proteins and small molecules that confer photosensitivity in *P. aeruginosa* biofilms as described here constitutes an initiative toward better understanding and treating host colonization by this bacterium.

## Materials and Methods

### Bacterial culture and colony morphology assay

Bacterial strains used in this study are listed in **Table S1**. For colony morphology assays, 1% tryptone + 1% agar was autoclaved, cooled to 60 °C, and 40 µg/mL Congo red (Alfa Aesar) and 20 µg/mL Coomassie blue (OmniPur, MilliporeSigma) dyes were added to the medium. The mixture was poured into 100 mm×100 mm×15 mm plates (LDP), 60 mL per plate, and left to cool and solidify overnight (14 to 24 h) at room temperature (25 °C). Overnight liquid cultures were grown in lysogeny broth (LB) at 37 °C with shaking at 250 rpm for 12 to 16 h. Overnight cultures were diluted 1:100 in fresh LB and subcultures were grown to mid-exponential phase (0.4 AU to 0.6 AU at OD_500_). Colony biofilms were seeded by spotting 10 µL of this bacterial subculture onto colony morphology medium. Spots were dried and incubated in a Percival CU-22LC9 incubator at 25 °C and with 90-100% humidity in the dark or under different light conditions (see below). Biofilm development was monitored by taking images at 24-h intervals using a Keyence VHX-1000 microscope, or by taking time-lapse movies in a custom-built movie recording chamber. Time lapse images were acquired at 15 min intervals by webcam (HD C920 and C930, Logitech) under 10 s white light panel (Porta-Trace, Gagne Inc.) illumination using a customized LabView (National Instruments) integration system.

### Colony morphology assay under light exposure

Colony morphology assay was prepared as described above and colony biofilms were incubated in a Percival CU-22LC9 incubator with a built-in lighting system. For all experiments, except for the light titration experiment (Fig. 1B) and light source experiment (Fig. 1C), white light exposure (bulbs: F17T8/TL841/ALTO, Philips) was calibrated to 16.5 µmol photons m^−2^ s^−1^ (7.35 µmol photons m^−2^ s^−1^ in the movie chamber). Light titration points (Fig. 1B) were 3.56 µmol photons m^−2^ s^−1^ (0.78 W m^−2^), 6.84 µmol photons m^−2^ s^−1^ (1.49 W m^−2^), 11.1 µmol photons m^−2^ s^−1^ (2.42 W m^−2^), and 16.6 µmol photons m^−2^ s^−1^ (3.62 W m^−2^), respectively. Light intensities were measured as photon flux density between 400 nm and 700 nm using an AMOUR-SL-125-PAR meter (Biospherical Instruments) for all light bulbs except the far-red light bulbs. For the light source experiment (Fig. 1C), white light intensity was set to 16.6 µmol photons m^−2^ s^−1^ (3.62 W m^−2^), blue light intensity (bulbs: Actinic Blue, F17T8, Coralife) was set to 13.8 µmol photons m^−2^ s^−1^ (3.83 W m^−2^), and red light intensity (bulbs: ELA-086, F17T8/IR-660, Percival Scientific) was set to 27.75 µmol photons m^−2^ s^−1^ (5.33 W m^−2^). Light intensities for peak wavelengths of all light bulbs were measured using a S130C photodiode power sensor (Thorlabs) with a PM100A power meter console (Thorlabs). For comparison between light bulbs, white light intensity (analyzed at 436, 545, and 612 nm) was measured as 2.66, 1.31, and 1.0 W m^−2^, respectively. Blue light intensity (analyzed at 422 nm) was measured as 0.7 W m^−2^, red light intensity (analyzed at 660 nm) was measured as 1.34 W m^−2^, and far-red light intensity (analyzed at 750 nm; bulbs: ELA-087, F17T8/IR-750, Percival Scientific) was measured as 1.04 W m^−2^.

### Cyclic di-GMP extraction and quantification from biofilms

Colony biofilms were spotted and grown as described above. Biofilms were scraped off the colony morphology agar plate at 24 h, 48 h, 72 h, 96 h, and 120 h. Scraped biofilms were transferred to a bead beater tube (19-649, Omni International) containing 1 mL of phosphate-buffered saline(PBS, pH 7.4) and 0.545 g of 1.4 mm zirconium oxide ceramic beads (19-645, Omni International). Biofilms were homogenized in a Bead Ruptor 12 Homogenizer (19-050, Omni International) for 99 s at high speed (5 m/s) at 4 °C. Eight hundred microliters of homogenized mixture were transferred to a cold microcentrifuge tube and centrifuged at 16,873×g for 60 s in a bench-top centrifuge (Model 5418, Eppendorf) at 4 °C. Cell pellets were resuspended in 250 µL of extraction buffer (methanol:acetonitrile:water (40:40:20) with 0.1 N formic acid) and incubated at −20 °C for 60 min,after which cell debris were pelleted by centrifugation at 21,130×g for 5 min. Two hundred microliters of supernatant were transferred to tubes containing 8 µL neutralization solution (15% NH4HCO3) and dried in a Vacufuge concentrator (Eppendorf) for 30 to 45 min. Samples were resuspended in Mobile Phase A (10 mM tributylamine + 15 mM Acetic acid] in Water:Methanol (97:3)) and shipped on dry ice to Mass Spectrometry Core Facility at Michigan State University. c-di-GMP concentrations of samples were quantified with a Quattro Premier XE LC/MS/MS using electrospray ionization analysis. The c-di-GMP concentration of each biofilm was normalized to the dried biomass of the respective sample, which was obtained by drying cell pellets in a Vacufuge concentrator (Eppendorf) and measuring biomass on an analytical scale (XS64, Mettler Toledo). Experiment was performed in biological triplicate.

### Colony morphology assay with exogenous phenazines

Colony morphology assay medium was prepared as described above and amended with 200µM phenazine-1-carboxylic acid (PCA, Apexmol), phenazine-1-carboxamide (PCN, Apexmol) or pyocyanin (PYO, Cayman Chemicals). The resulting medium was poured into 100 mm×100 mm×15 mm plates (LDP), 60 mL per plate, and left to cool and solidify overnight (14 to 24 h) at room temperature in the dark.

### Quantification of phenazine production

Colony morphology agar was prepared as described above and 4 mL of the mixture were poured into 35 mm×10 mm round Petri dishes (25373-041, VWR)and left to cool and solidify overnight (14 to 24 h) at 25 °C. Ten microliters of subcultures of WT PA14, grown as described above, were spotted onto a Petri dish containing colony morphology medium and grown for 96 h under either white light or dark conditions. Phenazines were harvested from each biofilm and the agar, upon which it was grown by methanol extraction: agar and biofilm were disrupted with a spatula and transferred to 5 mL of 100% methanol in a 15 mL polypropylene conical tube, and phenazines were extracted overnight in the dark, with constant agitation on a nutator. Three hundred microliters of extracted phenazine mixture were applied to a Spin-X column with 0.22 µm filter (VWR 29442-754) and centrifuged at 16,873×g for 5 min and 200 µL of flow-through was transferred to a sample vial. Concentrations of PCA, PCN and PYO were determined by high-performance liquid chromatography (1100 HPLC System, Agilent) using a method as described in (12, 21). Sample peaks were compared to peaks of pure phenazine standards for PCA, PCN and PYO, and the area under each peak was used to evaluate the concentration of each phenazine. For each condition, n=10.

### Construction of P*mexG*-*gfp* reporter strains

Transcriptional reporters of SoxR-driven *mexG* expression were constructed by transforming plasmid pLD2726 (58) into *Escherichia coli* S17-1 and moved into PA14 WT and ∆*phz* by conjugation as described in (59). The plasmid backbone was resolved out using FLP recombinase, introduced on plasmid pFLP2, and counterselected on sucrose plates as described in (59).

### Growth curve assay with phenazine oxidation state reporters

Overnight precultures of reporters (WT *attB*::P*mexG*-*gfp* and ∆*phz attB*::P*mexG*-*gfp*) were grown in 1% tryptone medium at 37 °C, shaking at 250 rpm, for 14 h. Overnight cultures were diluted 1:100 in fresh 1% tryptone medium and grown to mid-exponential phase (0.4 AU at OD_500_). Two identical assay plates were prepared simultaneously: cultures for the assay were started by inoculating 2 µL of subculture (in technical triplicate) into 200 µL of 1% tryptone per well in a black, flat-bottomed 96-well plate (655097, Greiner Bio); each strain was inoculated in biological triplicate. Both 96-well plates were placed in a 37 °C incubator with shaking at 300 rpm, one plate was placed under a dark box, while the other plate was exposed to white LED lights (22.8 µmol photons m^−2^ s^−1^). Cell density (OD_500_) and reporter expression (fluorescence at 480 nm excitation, 510 nm emission) were measured in a plate reader (BioTek) every hour for the first six hours and every two hours for the next 18 hours. Analysis was performed by normalizing each fluorescence value to the corresponding growth value, averaging the technical replicates, followed by averaging the biological replicates.

### Statistical analysis

All statistical analyses were performed for at least three biological replicates per condition. *p*-values were determined using unpaired two-tailed *t*-test, in most cases by comparing data from light condition to data from dark condition (\**p*≤0.05, \*\**p*≤0.01, \*\*\*\**p*≤0.0001, n.s. = not significant). For light titration experiment (Fig. 1B), *p*-values were determined, comparing data from light level of interest to data from next highest light level (\*\*\*\**p*≤0.0001, n.s. = not significant).

## Acknowledgements

We thank Yasuhiko Irie for discussion and experimental feedback. We thank Andreas Hartel, William Stoy and Joaquim Goes for technical assistance with the light measurements. We thank Lijun Chen at the Michigan State University RTSF Mass Spectrometry Core Facility for help with the c-di-GMP analysis. This work was supported by NIH/NIAID grant R01AI103369 and an NSF CAREER award to L.E.P.D.

## References

1. Purcell EB, Crosson S. 2008. Photoregulation in prokaryotes. Curr Opin Microbiol 11:168–178.

2. Gomelsky M, Hoff WD. 2011. Light helps bacteria make important lifestyle decisions. Trends Microbiol 19:441–448.

3. Bonomi HR, Toum L, Sycz G, Sieira R, Toscani AM, Gudesblat GE, Leskow FC, Goldbaum FA, Vojnov AA, Malamud F. 2016. *Xanthomonas campestris* attenuates virulence by sensing light through a bacteriophytochrome photoreceptor. EMBO Rep 17:1565–1577.

4. Purcell EB, Siegal-Gaskins D, Rawling DC, Fiebig A, Crosson S. 2007. A photosensory two-component system regulates bacterial cell attachment. Proceedings of the National Academy of Sciences.

5. Pané-Farré J, Quin MB, Lewis RJ, Marles-Wright J. 2017. Structure and Function of the Stressosome Signalling Hub. Subcell Biochem 83:1–41.

6. Kim H-S, Willett JW, Jain-Gupta N, Fiebig A, Crosson S. 2014. The *Brucella abortus* virulence regulator, LovhK, is a sensor kinase in the general stress response signalling pathway. Mol Microbiol 94:913–925.

7. Agostoni M, Koestler BJ, Waters CM, Williams BL, Montgomery BL. 2013. Occurrence of cyclic di-GMP-modulating output domains in cyanobacteria: an illuminating perspective. MBio 4.

8. Herrou J, Crosson S. 2011. Function, structure and mechanism of bacterial photosensory LOV proteins. Nat Rev Microbiol 9:713–723.

9. Tarutina M, Ryjenkov DA, Gomelsky M. 2006. An unorthodox bacteriophytochrome from *Rhodobacter sphaeroides* involved in turnover of the second messenger c-di-GMP. J Biol Chem 281:34751–34758.

10. Barends TRM, Hartmann E, Griese JJ, Beitlich T, Kirienko NV, Ryjenkov DA, Reinstein J, Shoeman RL, Gomelsky M, Schlichting I. 2009. Structure and mechanism of a bacterial light-regulated cyclic nucleotide phosphodiesterase. Nature 459:1015–1018.

11. Tschowri N, Busse S, Hengge R. 2009. The BLUF-EAL protein YcgF acts as a direct anti-repressor in a blue-light response of *Escherichia coli*. Genes Dev 23:522–534.

12. Jo J, Cortez KL, Cornell WC, Price-Whelan A, Dietrich LE. 2017. An orphan cbb3-type cytochrome oxidase subunit supports *Pseudomonas aeruginosa* biofilm growth and virulence. Elife 6.

13. Okegbe C, Fields BL, Cole SJ, Beierschmitt C, Morgan CJ, Price-Whelan A, Stewart RC, Lee VT, Dietrich LEP. 2017. Electron-shuttling antibiotics structure bacterial communities by modulating cellular levels of c-di-GMP. Proc Natl Acad Sci U S A 114:E5236–E5245.

14. Price-Whelan A, Dietrich LEP, Newman DK. 2007. Pyocyanin alters redox homeostasis and carbon flux through central metabolic pathways in *Pseudomonas aeruginosa* PA14. J Bacteriol 189:6372–6381.

15. Dietrich LEP, Okegbe C, Price-Whelan A, Sakhtah H, Hunter RC, Newman DK. 2013. Bacterial community morphogenesis is intimately linked to the intracellular redox state. J Bacteriol 195:1371–1380.

16. Recinos DA, Sekedat MD, Hernandez A, Cohen TS, Sakhtah H, Prince AS, Price-Whelan A, Dietrich LEP. 2012. Redundant phenazine operons in *Pseudomonas aeruginosa* exhibit environment-dependent expression and differential roles in pathogenicity. Proc Natl Acad Sci U S A 109:19420–19425.

17. Friedman L, Kolter R. 2004. Genes involved in matrix formation in *Pseudomonas aeruginosa* PA14 biofilms. Mol Microbiol 51:675–690.

18. Pfeifroth U, Sanchez-Lorenzo A, Manara V, Trentmann J, Hollmann R. 2018. Trends and variability of surface solar radiation in Europe based on surface-and satellite-based data records. J Geophys Res D: Atmos 123:1735–1754.

19. Lee VT, Matewish JM, Kessler JL. 2007. A cyclic-di-GMP receptor required for bacterial exopolysaccharide production. Molecular Microbiology.

20. Hickman JW, Harwood CS. 2008. Identification of FleQ from *Pseudomonas aeruginosa* as a c-di-GMP-responsive transcription factor. Mol Microbiol.

21. Sakhtah H, Koyama L, Zhang Y, Morales DK, Fields BL, Price-Whelan A, Hogan DA, Shepard K, Dietrich LEP. 2016. The *Pseudomonas aeruginosa* efflux pump MexGHI-OpmD transports a natural phenazine that controls gene expression and biofilm development. Proc Natl Acad Sci U S A 113:E3538–47.

22. Flood ME, Herbert RB, Holliman FG. 1972. Pigments of *Pseudomonas* species. Part V. Biosynthesis of pyocyanin and the pigments of *Ps. aureofaciens*. J Chem Soc Perkin 10:622–626.

23. Hansford GS, Holliman FG, Herbert RB. 1972. Pigments of *Pseudomonas* species. Part IV. *in vitro* and *in vivo* Conversion of 5-methylphenazinium-1-carboxylate into aeruginosin A. J Chem Soc Perkin 1 103–105.

24. Abu EA, Su S, Sallans L, Boissy RE, Greatens A, Heineman WR, Hassett DJ. 2013. Cyclic voltammetric, fluorescence and biological analysis of purified aeruginosin A, a secreted red pigment of *Pseudomonas aeruginosa* PAO1. Microbiology.

25. Mavrodi DV, Bonsall RF, Delaney SM, Soule MJ, Phillips G, Thomashow LS. 2001. Functional analysis of genes for biosynthesis of pyocyanin and phenazine-1-carboxamide from *Pseudomonas aeruginosa* PAO1. J Bacteriol 183:6454–6465.

26. Parsons JF, Greenhagen BT, Shi K, Calabrese K, Robinson H, Ladner JE. 2007. Structural and functional analysis of the pyocyanin biosynthetic protein PhzM from *Pseudomonas aeruginosa*. Biochemistry 46:1821–1828.

27. Greenhagen BT, Shi K, Robinson H, Gamage S, Bera AK, Ladner JE, Parsons JF. 2008. Crystal structure of the pyocyanin biosynthetic protein PhzS. Biochemistry 47:5281–5289.

28. Dietrich LEP, Teal TK, Price-Whelan A, Newman DK. 2008. Redox-active antibiotics control gene expression and community behavior in divergent bacteria. Science 321:1203–1206.

29. Dietrich LEP, Price-Whelan A, Petersen A, Whiteley M, Newman DK. 2006. The phenazine pyocyanin is a terminal signalling factor in the quorum sensing network of *Pseudomonas aeruginosa*. Mol Microbiol 61:1308–1321.

30. Gu M, Imlay JA. 2011. The SoxRS response of Escherichia coli is directly activated by redox-cycling drugs rather than by superoxide. Mol Microbiol 79:1136.

31. Mackow N, Dietrich L, Chander M. 2013. Species-specific residues calibrate SoxR sensitivity to redox-active molecules. Molecular Microbiology.

32. Römling U, Galperin MY, Gomelsky M. 2013. Cyclic di-GMP: the first 25 years of a universal bacterial second messenger. Microbiol Mol Biol Rev 77:1–52.

33. Ha D-G, O’Toole GA. 2015. c-di-GMP and its Effects on Biofilm Formation and Dispersion: a *Pseudomonas Aeruginosa* Review. Microbiol Spectr 3:MB-0003-2014.

34. Dayton H, Smiley MK, Forouhar F, Harrison JJ, Price-Whelan A, Dietrich LE. in press. Sensory domains that control cyclic di-GMP-modulating proteins: a critical frontier in bacterial signal transduction. Springer.

35. Christen M, Kulasekara HD, Christen B, Kulasekara BR, Hoffman LR, Miller SI. 2010. Asymmetrical distribution of the second messenger c-di-GMP upon bacterial cell division. Science 328:1295–1297.

36. Henry JT, Crosson S. 2011. Ligand-binding PAS domains in a genomic, cellular, and structural context. Annu Rev Microbiol 65:261–286.

37. Garcia D, Watts KJ, Johnson MS, Taylor BL. 2016. Delineating PAS-HAMP interaction surfaces and signalling-associated changes in the aerotaxis receptor Aer. Mol Microbiol 100:156–172.

38. Gong W, Hao B, Mansy SS, Gonzalez G, Gilles-Gonzalez MA, Chan MK. 1998. Structure of a biological oxygen sensor: a new mechanism for heme-driven signal transduction. Proc Natl Acad Sci U S A 95:15177–15182.

39. Taylor BL, Zhulin IB. 1999. PAS domains: internal sensors of oxygen, redox potential, and light. Microbiol Mol Biol Rev 63:479–506.

40. Taylor BL, Rebbapragada A, Johnson MS. 2001. The FAD-PAS domain as a sensor for behavioral responses in *Escherichia coli*. Antioxid Redox Signal 3:867–879.

41. Kuchma SL, Brothers KM, Merritt JH, Liberati NT, Ausubel FM, O’Toole GA. 2007. BifA, a cyclic-Di-GMP phosphodiesterase, inversely regulates biofilm formation and swarming motility by *Pseudomonas aeruginosa* PA14. J Bacteriol 189:8165–8178.

42. Roy AB, Petrova OE, Sauer K. 2012. The phosphodiesterase DipA (PA5017) is essential for *Pseudomonas aeruginosa* biofilm dispersion. J Bacteriol 194:2904–2915.

43. Auldridge ME, Forest KT. 2011. Bacterial phytochromes: more than meets the light. Crit Rev Biochem Mol Biol 46:67–88.

44. Kopka B, Magerl K, Savitsky A, Davari MD, Röllen K, Bocola M, Dick B, Schwaneberg U, Jaeger K-E, Krauss U. 2017. Electron transfer pathways in a light, oxygen, voltage (LOV) protein devoid of the photoactive cysteine. Sci Rep 7:13346.

45. Losi A, Gärtner W. 2011. Old chromophores, new photoactivation paradigms, trendy applications: flavins in blue light-sensing photoreceptors. Photochem Photobiol 87:491–510.

46. Glasser NR, Wang BX, Hoy JA, Newman DK. 2017. The pyruvate and α-ketoglutarate dehydrogenase complexes of *Pseudomonas aeruginosa* catalyze pyocyanin and phenazine-1-carboxylic acid reduction via the subunit dihydrolipoamide dehydrogenase. J Biol Chem 292:5593–5607.

47. Bury A, Hellingwerf KJ. 2014. On the *in vivo* redox state of flavin-containing photosensory receptor proteins. Methods Mol Biol 1146:177–190.

48. Purcell EB, McDonald CA, Palfey BA, Crosson S. 2010. An analysis of the solution structure and signaling mechanism of LovK, a sensor histidine kinase integrating light and redox signals. Biochemistry 49:6761–6770.

49. Katayama B, Ozawa T, Morimoto K, Awazu K, Ito N, Honda N, Oiso N, Tsuruta D. 2018. Enhanced sterilization and healing of cutaneous pseudomonas infection using 5-aminolevulinic acid as a photosensitizer with 410-nm LED light. J Dermatol Sci 90:323–331.

50. Leanse LG, Dong P-T, Goh XS, Lu M, Cheng J-X, Hooper DC, Dai T. 2019. Quinine Enhances Photo-Inactivation of Gram-Negative Bacteria. J Infect Dis.

51. Hendiani S, Rybtke ML, Tolker-Nielsen T, Kashef N. 2019. Sub-lethal antimicrobial photodynamic inactivation affects *Pseudomonas aeruginosa* PAO1 quorum sensing and cyclic di-GMP regulatory systems. Photodiagnosis Photodyn Ther 27:467–473.

52. Baron SS, Rowe JJ. 1981. Antibiotic action of pyocyanin. Antimicrob Agents Chemother 20:814–820.

53. Curtis AM, Bellet MM, Sassone-Corsi P, O’Neill LAJ. 2014. Circadian clock proteins and immunity. Immunity 40:178–186.

54. Swartz TE, Tseng T-S, Frederickson MA, Paris G, Comerci DJ, Rajashekara G, Kim J-G, Mudgett MB, Splitter GA, Ugalde RA, Goldbaum FA, Briggs WR, Bogomolni RA. 2007. Blue-light-activated histidine kinases: two-component sensors in bacteria. Science 317:1090–1093.

55. Santamaría-Hernando S, Rodríguez-Herva JJ, Martínez-García PM, Río-Álvarez I, González-Melendi P, Zamorano J, Tapia C, Rodríguez-Palenzuela P, López-Solanilla E. 2018. *Pseudomonas syringae* pv. tomato exploits light signals to optimize virulence and colonization of leaves. Environ Microbiol 20:4261–4280.

56. Idnurm A, Crosson S. 2009. The photobiology of microbial pathogenesis. PLoS Pathog 5:e1000470.

57. Mukherjee S, Jemielita M, Stergioula V, Tikhonov M. 2019. Photo sensing and quorum sensing are integrated to control bacterial group behaviors. bioRxiv.

58. Sporer AJ, Beierschmitt C, Bendebury A, Zink KE, Price-Whelan A, Buzzeo MC, Sanchez LM, Dietrich LEP. 2018. *Pseudomonas aeruginosa* PumA acts on an endogenous phenazine to promote self-resistance. Microbiology 164:790–800.

59. Lin Y-C, Sekedat MD, Cornell WC, Silva GM, Okegbe C, Price-Whelan A, Vogel C, Dietrich LEP. 2018. Phenazines Regulate Nap-Dependent Denitrification in *Pseudomonas aeruginosa* Biofilms. J Bacteriol 200.

